# Female and male *Sirex noctilio* use age and size to select a mate

**DOI:** 10.1101/2021.02.14.431120

**Authors:** Joséphine Queffelec, Jeremy D. Allison, Bernard Slippers, Jaco M. Greeff

## Abstract

While male mate choice in insects is a widely accepted concept, there is still limited evidence showing that lek formation is compatible with the evolution of male mate choice. In the woodwasp *Sirex noctilio*, males form leks that are used by females to select a mate. However, males have been observed to ignore certain females, suggesting the presence of male mate choice despite the presence of a lek mating system. In this study we demonstrate that males only attempt to mate with certain females. To understand the criteria used by males and females to select a mate, we also tested the effect of age, size, and male to female size ratio on the number of mating attempts made by males and on female receptivity. We demonstrate that size and age play a role in both male and female mate choice. Our results suggest that males must reach sexual maturity after emergence and are neither receptive nor attractive to females during the first few days of their lives. We also show that older females become less attractive to males, suggesting that female *S. noctilio* switch to a strict host location phase sometime after emergence. Our results show that male and female size, and the ratio between them, play a role in mate choice. While larger males are more motivated to mate, their large size can physically prevent them from mating with small females. Small females are also more attractive and more receptive to males, consistent with the presence of convenience polyandry in *S. noctilio*.

---

To facilitate mate location, some insect species have evolved lekking behaviours (Alcock 1987; Shelly and Whittier 1997). Leks are defined as male-biased aggregations formed for mating, where sperm is the only resource acquired by visiting females (Höglund and Alatalo 1995). These male aggregations are advantageous for both sexes because they can reduce predation risks, increase the probability of mating, facilitate mate location or allow individuals to rapidly assess and compare mate quality (Höglund and Alatalo 1995; Field et al. 2002). Within lek forming species, females use a variety of olfactory, visual and auditory cues and signals to evaluate male quality. These cues and signals deliver information about male life history, morphology, behaviour and within lek hierarchy (Lederhouse 1982; Jang and Greenfield 1996; Kaspi et al. 2000; Cooley and Marshall 2004; Segura et al. 2007; Shelly et al. 2007; Izzo and Tibbetts 2012).

In non-lekking species, three factors can lead to the evolution of male mate choice; variance in female quality (Nandy et al. 2012), high cost of mating due to time or energy constraints (Dewsbury 1982), and low cost of searching and assessment of females (Johnstone et al. 1996). In lekking species however, it is often believed that the extreme competition between males would outweigh the benefits of male mate choice and prevent its evolution. Despite these assumptions, a few studies have demonstrated the presence of male mate choice in some lekking species (Sæther et al. 2001; Werner and Lotem 2003; Shelly et al. 2012).

The woodwasp *Sirex noctilio* Fabricus (Hymenoptera: Siricidae) lays eggs in the wood of various pine species where the larvae bore galleries and feed on the wood before emerging as adults (Slippers et al. 2012). While *S. noctilio* originates from Eurasia (Spradbery and Kirk 1978), it has been accidently introduced in many countries of the Southern Hemisphere and causes large economic losses by killing planted pine trees (Hurley et al. 2007). In more recent years, the wasp has been introduced into countries of the Northern Hemisphere (i.e. the USA, Canada and China (Hoebek et al. 2005; Li et al. 2015)).

After emerging from the wood,male *S. noctilio* fly to the canopy to aggregate and form leks (Madden 1988). The females join these leks, mate with multiple males and fly away to find a host to lay eggs. While we understand how both sexes encounter each other by meeting at the top of pine trees, it is still unclear how pairs are formed. Is mate choice exhibited by one or both sexes? What traits are used by males or females to evaluate the quality of a potential mate? Is there a hierarchy within the lek that dictates who mates with the female?

In insects, body size is a trait commonly used by males to choose a mate because it is a good indicator of fecundity (Bonduriansky 2001). In *S. noctilio*, both males and females exhibit marked size variation with size ranging from 12 to 34 mm in females and from 9.3 to 34.9 mm in males (Neumann et al. 1987). For this reason, we hypothesize that size may play a role in mate choice in *S. noctilio*. Furthermore, it has been observed in captivity that large males often struggle to bend their abdomen to fertilize small females (Caetano and Hajek 2017). We hypothesize that the size ratio between a male and a female could also influence mating success in *S. noctilio*.

Age is considered a simple and efficient way to display “good genes” to a potential mate (Brooks and Kemp 2001). Male *S. noctilio* usually emerge a few days to a few weeks before females (Rawlings 1948; Morgan and Stewart 1966; Haavik et al. 2013). While they are waiting for the females to emerge, males are exposed to predators, parasites and to abiotic stresses that can reduce their life span. We hypothesize that female *S. noctilio* consider age when choosing a mate and preferentially mate with older males. On the other hand, because females have a very short life span of approximately 5 days (Neumann et al. 1987) and do not feed as adults (Morgan and Stewart 1966; Taylor 1981), we hypothesize that males would benefit from mating preferentially with young females that have more time to use their sperm.

We determine if size, age and male to female size ratio affect the motivation of males and females to mate and their mating success.

## MATERIAL AND METHODS

### Rearing of S. noctilio

Logs of *Pinus patula* and *P. radiata* were collected in the South African provinces of the Western Cape and KwaZulu-Natal between September 2018 and January 2019. The logs were brought to the Forestry and Agricultural Biotechnology Institute Biocontrol Centre of the University of Pretoria and placed in emergence cages and maintained at 23± 10°C; 12L:12D.

*Sirex noctilio* adults were collected from the emergence cages daily between 0700 and 0900, placed in individual plastic containers (12 cm x 12 cm x 5 cm) and kept at 23°C with a photoperiod of 12L:12D until they were used for bioassays. The bottom of the boxes were covered in pieces of pine bark because *S. noctilio* adults struggle to gain traction on the smooth surface of plastic containers. The pine bark was sprayed with distilled water every two days to prevent dehydration of the wasps. Those conditions allowed us to reduce wasp stress. Under these conditions, males lived up to 20 days and females up to 12 days, indicating that the wasps were not stressed and could be used for behavioural tests.

During collection of adults, if emergence cages contained more than one sex, the individuals in that cage were discarded to ensure that all individuals used for behavioural tests were naïve.

### Bioassays

*A* total of 21 bioassays were conducted. The bioassays were conducted in a wooden cage (31 cm x 32 cm x 42 cm) with one glass screen wall. All walls, except for the glass screen, were covered with paper and changed after each bioassay to avoid contamination through the deposition of contact and volatile pheromones. Furthermore, the glass screen was cleaned with 70% ethanol after each bioassay. On the floor of the cage, a grid was drawn to facilitate scoring the location of individuals.

The cage was placed outside on a table in the shade. *Sirex noctilio* adults are more likely to mate at higher temperatures (Caetano and Hajek 2017). To guarantee that low temperatures would not interfere with mating success, all bioassays were conducted between 0800 and 1230 on days with a cloud cover below 50%. The temperature varied between 22°C and 32°C.

One female was first introduced into the cage and kept under a small plastic cage (10 cm x 10 cm x 5 cm). This allowed the female to recover from the stress of handling. The placement of the small plastic cage in the cage was standardized across bioassays. Eight males were then randomly selected, and a dot of oil-based paint (Dala oil paint) was painted on the back of the thorax using a different colour for each male. The males where then introduced into the cage. During the first few minutes after being introduced into the cage, the males usually exhibit aggressive behaviours towards females and each other. To prevent injuries to the female, the small plastic cage isolating her was only removed 5 minutes after the introduction of the males. The bioassays lasted 20 min starting from the removal of the small plastic cage.

### Data capture

Before being placed into the cage, the age of each male and female were recorded. All bioassays were recorded using a video camera (Canon LEGRIA HF G26). Oral description of the location of the males in the cage was done continuously as the dots of paints on the males were not always visible on the video. The data extracted from the video were the number of contacts between the males and each female, the number of mating attempts made by each male towards the female and the number of matings between the female and each male.

When placed in a cage with a female, males have rarely been observed to walk towards the female in a straight line. Males must come into contact with a female to detect her contact pheromones (Dolezal 1967; Böröczky et al. 2009). It is only after a physical contact between the antennae of the male and the body of the female that a male will initiate mating. As a result, contact between a male and a female was defined as a contact between the male antennae and any part of the female body. Only the bioassays during which each male came into contact with the female at least once were used.

To initiate mating and fertilize the female, a male must grasp the female with his front legs, bend his abdomen to reach underneath the female and open his claspers. The claspers are located on the sides of the male genitalia and allow the males to lock themselves on the female’s genitalia and to prevent the mating from being interrupted before fertilization. A mating attempt was defined as a male bending his abdomen forward and opening his claspers within one second of a contact with a female.

On average, we observe that a mating in *S. noctilio* lasts between 20 seconds and one minute where the male is locked onto the female with its claspers. Therefore, a mating was defined as a male being visibly locked to a female for more than 20 seconds.

After each bioassay, all individuals were placed in 90% ethanol and later the width of the pronotum (mm) was measured as a proxy for size (Madden 1981; Haavik et al. 2016). The measurements were taken using a pair of digital callipers.

### Statistical analysis of the male motivation to mate and mating success

The number of times a male mates depends on its motivation to mate and on the response of the female, both of which can be influenced by the age and size of the male. We used two response variables; male motivation to mate, quantified as the number of attempts exhibited by a male, and male mating success, measured by the number of times a male mated.

Eight males were placed in one cage with one female. While this strategy recreates lek conditions, when *n* males are caged with a female, each of the male observations are interdependent and provide only one data point. To test a single variable, *n*-1 males must be kept identical while the *n*^th^ male must be varied for this trait. To estimate the role of a variable using standard statistics is logistically arduous and to estimate the effect of more than one variable becomes logistically impossible. To circumvent this problem, we used an approach advocated by Hilborn and Mangel (1997) that relies on likelihood. We did the same analysis on male motivation to mate and male mating success. The method is summarized below using the example of male mating success, but the same applies to male motivation to mate.

We calculated the likelihoods for a series of nested models that predict the number of matings for each male. We then searched for values of the variables that maximized the likelihood of our observed data given each model. Finally, we compared the models using the values of negative log-likelihood, AICc (Hurvich and Tsai 1989) and BIC (Schwarz 1978). When a model was nested in another (i.e. the parameters estimated in the smaller model are a subset of the parameters estimated in the larger model), these models could be compared using a likelihood ratio test. For the best model we used a 95% confidence interval and a likelihood ratio test for each variable by comparing the model to the maximum likelihood.

We considered eight variables in a family of nested models (Figure 1). Given values of all variables and a male *i*’s age (*A*), pronotum width (*S*_m_) and his relative size (*S*_r_) compared to the size of the female in his cage (*S*_f_) (*S*_r_=*S*_m_/*S*_f_), we calculate a score for each male as follows:

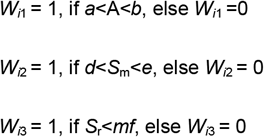

**Figure 1:**
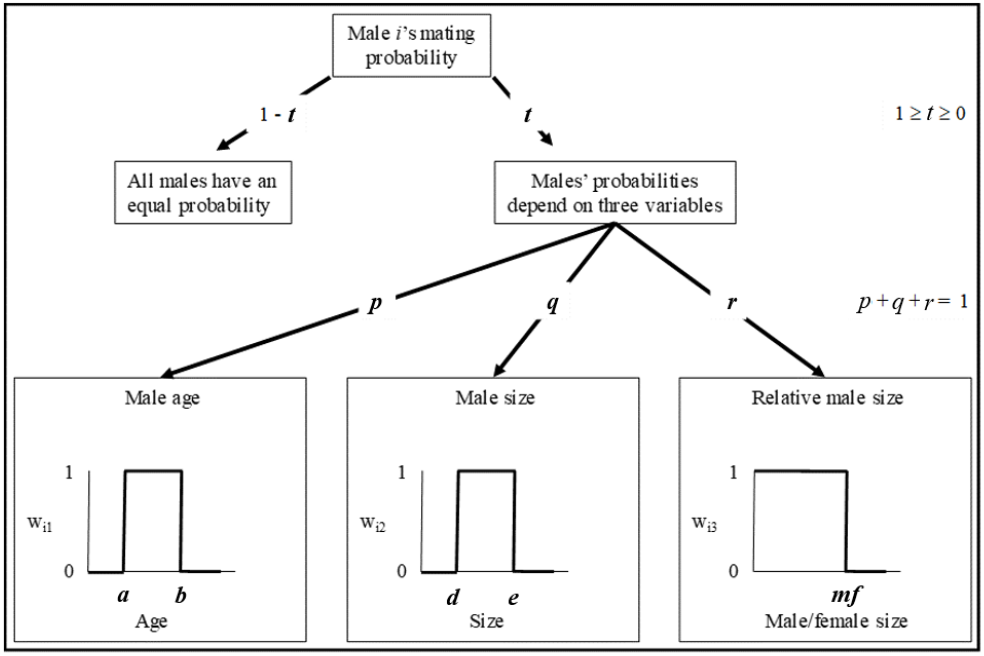
Diagram showing how the probabilities of males’ mating attempts and success are calculated using nine variables. Because *p* + *q* + *r* = 1, eight variables define the full model.

With *mf* = male/female size ratio. These scores are combined in an overall score of the *i*th male as:

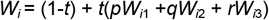

The *W* value of each male is used as the relative probabilities in a multinominal distribution, which is compared to the actual mating distribution observed in the cage. This calculation is the likelihood of the data given the model variables. The log-likelihoods of each cage are then added to obtain an overall likelihood of the model explaining the data. By setting specific variables equal to the minimum and maximum values observed for each variable, we achieve a nested series of models that can be compared to each other.

We wrote functions in R (R Core Team 2017) to calculate how the likelihood varies for one variable while all the other variables are fixed. These functions consider a specific number of equal steps in variable space for the one variable given a lower and upper limit. We worked through a series of such functions, one for each variable, iteratively to find the variable values that maximized the likelihood of each model. Once the maximum likelihood was obtained, BIC, AICc and negative log-likelihood could be calculated to compare the models.

Because the models are nested (e.g. the parameters estimated in model G15 are a subset of the parameters estimated in model G7), we can use a likelihood ratio test to investigate if the likelihood differences are significant. If L{G15} and L{G7} are the negative log-likelihood of model G15 and G7 respectively, and if model G15 is nested in model G7, then, if 2(L{G15}-L{G7})>Chi-squared, with the degrees of freedom equal to the difference in the number of parameters between the models, there is a significant difference between model G15 and model G7.

The use of a likelihood-based model prevents the use of data points with zero observations. When a female did not mate with any of the males in her cage, her bioassay was removed from the male mating dataset, this resulted in the removal of six bioassays. Furthermore, during the data exploration phase, it became evident that one data point was biasing the results due to a single male making 15 mating attempts which was significantly higher than the average 1.15 attempts per male. This bioassay was removed from the dataset for the analysis of the number of attempts by males.

### Statistical analysis of female attractiveness and female receptivity

The mating success of a female depends on the number of males that attempt mating with her and on the number of times she will accept a male. Both types of events can be influenced by the age and size of both sexes. Therefore, we used two response variables; female attractiveness defined as the number of attempts a female receives from the males in her cage, and female receptivity defined as the proportion of attempts that lead to a mating.

Because each female was independently tested by introducing one female per cage, female attractiveness and receptivity could be analysed using generalized linear mixed models. Because female attractiveness is a count variable, we used a generalized linear mixed model with Poisson errors. Female receptivity is a binomial variable that includes the number of attempts by a male leading to a mating (success) and the number of attempts by a male leading to a rejection of the male by the female (failure). For the analysis of female receptivity, we used a generalized linear mixed model with binomial errors.

We predicted female attractiveness (called model I) and female receptivity (called model J) by starting with models predicting female attractiveness and female receptivity as functions of female pronotum width (mm), female age (days), male pronotum width (mm), male age (days), male to female pronotum width ratio and the interactions between female age and female pronotum width and between male age and male pronotum width. The female number was added to both models I and J as a random effect. For model selection a stepwise backward elimination process was performed, using deviance analysis with an F-test until a minimal adequate model was obtained.

## RESULTS

### Female mating success and male mating skew

The average number of matings per female was *X* +SE = 1.86 + 0.48, *N* = 21. Of the 15 females that mated (71.4% of the females), 66.7% of them mated more than once. On average

*X* + SE = 19.7 + 12.5%, *N* = 21 of contacts between a male and a female led to an attempt by the male. Of the 168 males used in the bioassays, 28 of them obtained all of the 39 matings recorded.

### Male motivation to mate

Table 1 gives the negative log-likelihood, AICc and BIC for a set of nested models illustrated in Figure 1. Each row gives the optimized model parameters for the model. We can use the AICc and BIC to select the best models as these indices consider the sample size and the number of parameters estimated (*k*). Lower values of the negative log-likelihood, AICc and BIC indicate better-fitted models.

**Table 1:**
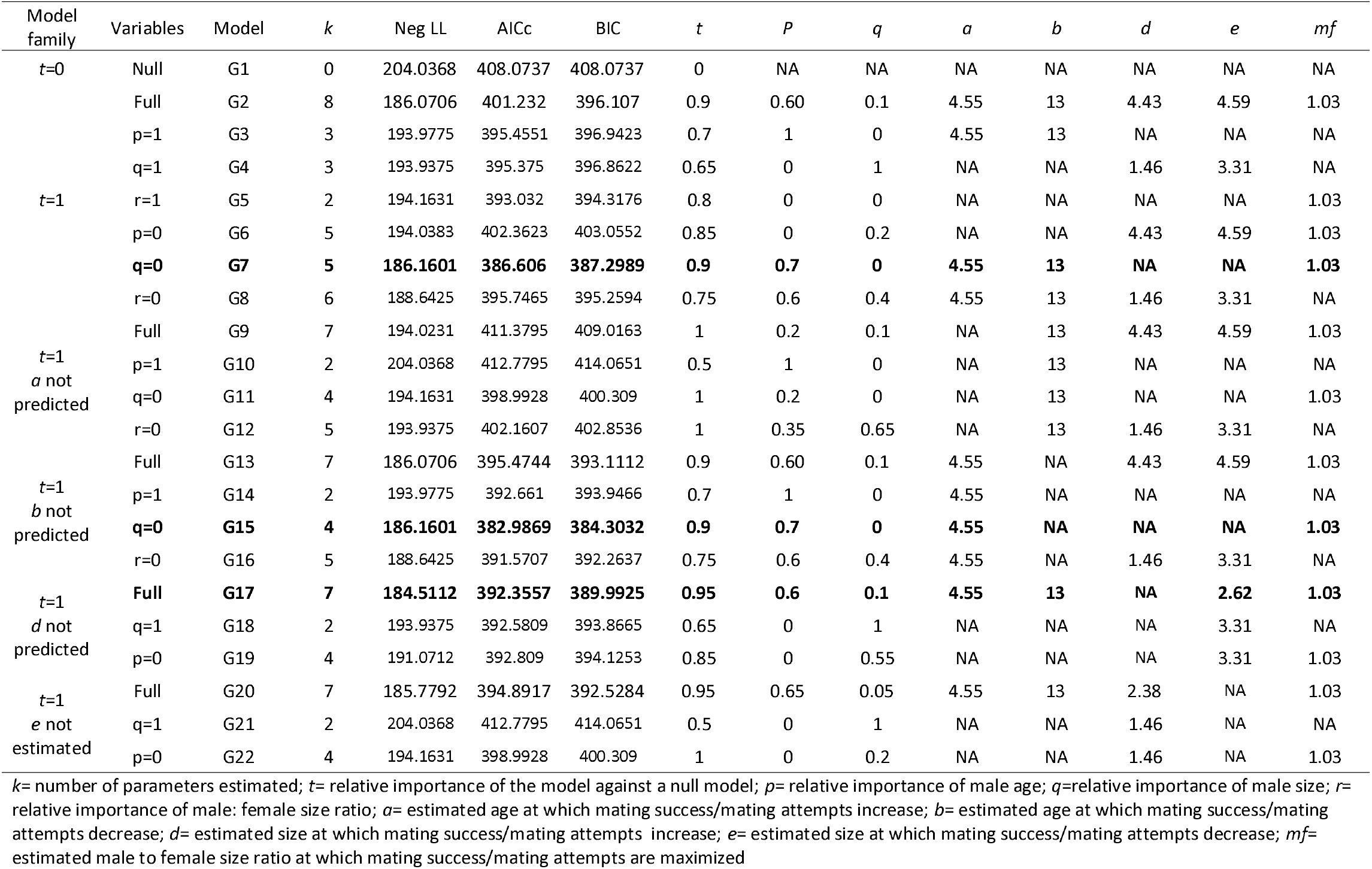
Summary of the maximum likelihood model parameters for the nested models when predicting the number of mating attempts by each male.

Considering the AICc and the BIC, model G15 is the best, followed by model G7 (Table 1). Both models G15 and G7 suggested male age (70%) and male to female pronotum width ratio (30%) played a role in predicting the number of attempts made by males. The number of attempts increased in males older than four days old. The third-best model is G17 where male age (60%), male pronotum width (10%) and male to female pronotum width ratio (30%) played a role in predicting the number of attempts made by males. The number of attempts increased in males older than four days old and decreased in males with a pronotum width of over 2.62 mm. The average male pronotum width was *X* + SE = 3.11 + 0.96 mm, *N* = 168. Sixty-three percent of the males used in the bioassays had a pronotum width over 2.62 mm. All three models G15, G7 and G17 also predicted that the number of attempts decreased in males larger than 103% the size of the female.

The likelihood ratio test found no significant difference between models G15 and G7 (*P* > 0.05). Because model G17 was not nested in model G15 nor G7, it could not be compared through a likelihood ratio test. However, model G17 had a higher AICc and BIC compared to models G7 and G15, suggesting that it is not as good.

Figure 2 shows the fit of model G7 when *t, p, a, b* and *mf* are allowed to vary. The variation around the optimal *t* was tight and the variation around the optimal *p* was broad (0.45 < *P* < 0.83). The effect of variation in *a* was erratic, and not well behaved but only one peak had a higher likelihood (*a* = 4.55). The likelihood of *b* was also bimodal with a second peak being within the 95% confidence interval of the maximum likelihood. The distribution of *mf* was unimodal with a fairly tight variation around the optimal switch value.

**Figure 2:**
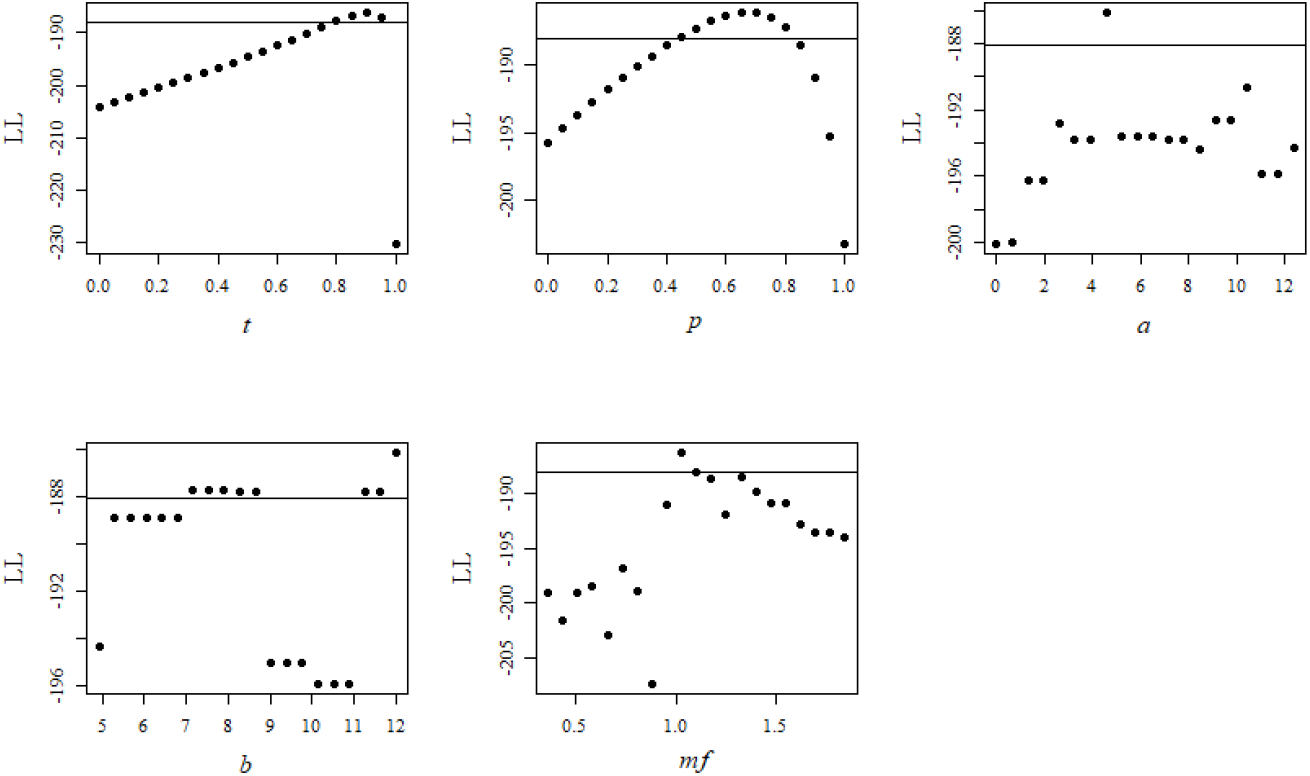
Variation in log-likelihood when *t, p, a, b* and *mf* are not fixed for model G7. The horizontal line is set at LL = -(negative log-likelihood + Chi-squared (*P* = 0.95, df = 1) /2). This value of LL represents a significant difference in log-likelihood between the point under and over the line.

### Male mating success

Based on the AICc and the BIC, model H14 seemed the best (Table 2). Model H14 suggested that only male age played a role in male mating success, with mating success increasing in males older than four days old. Model H15 suggested that male age was the most important predictor (85%) of male mating success, with males older than four days mating more. In this model, male to female pronotum width ratio also played a role (15%), with males smaller than 82% of the female size mating more often. The third-best model, H3, was similar to H14 in that only age played a role and no decrease in male mating success was found in older males.

**Table 2:**
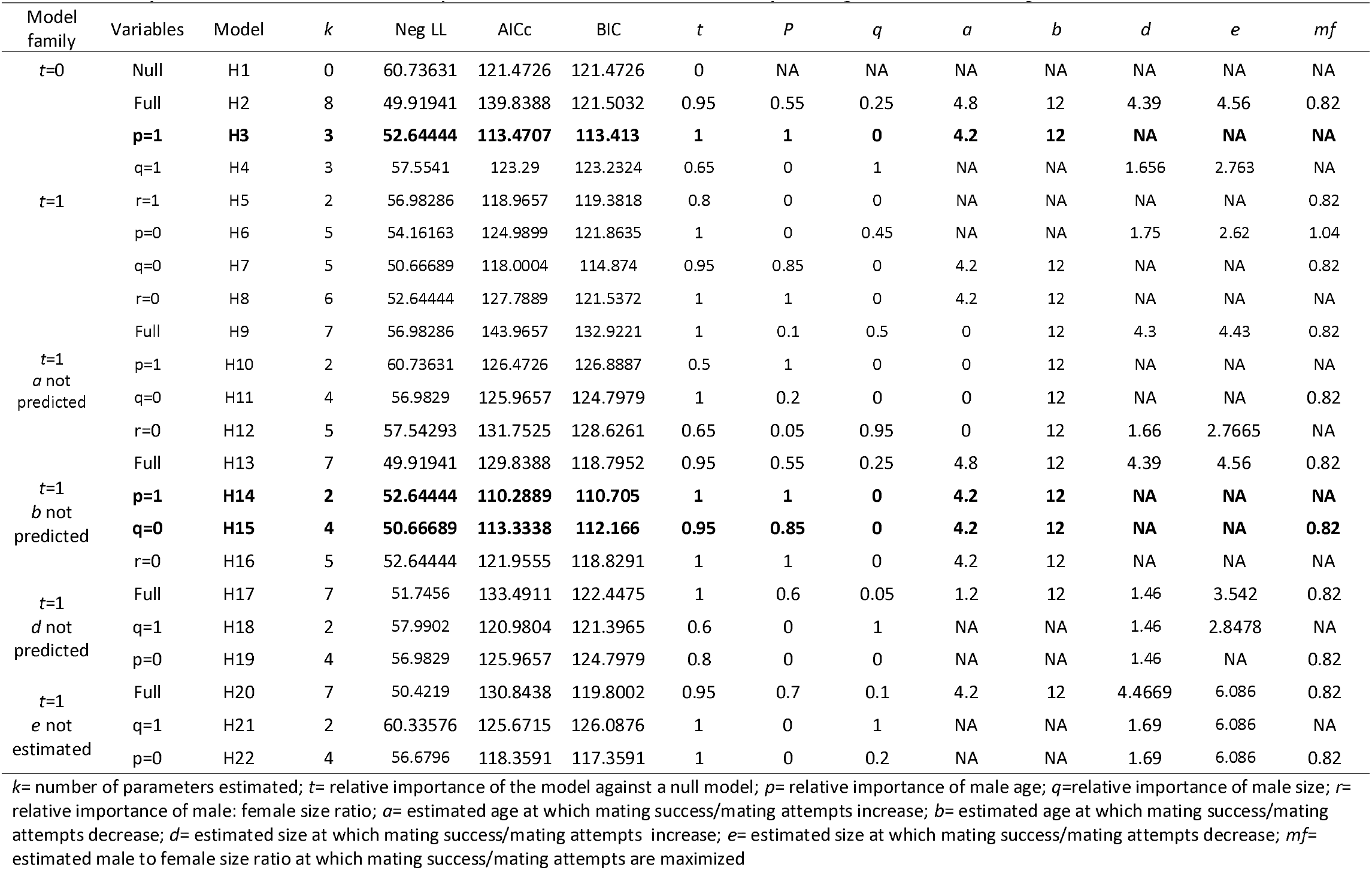
Summary of the maximum likelihood model parameters for the nested models when predicting the number of matings between each male and the female

The likelihood ratio test found no significant difference between models H14 and H15 *(P* > 0.05). However, model H14 was significantly better than model H3 (*P* < 0.05). Because models H15 and H3 were not nested, they could not be compared through a likelihood ratio test. However, model H15 had a lower AICc and BIC than model H3 and was thus better.

In Figure 3, the variation around the optimal *t* was tight and the variation around the optimal *p* was large (0.425 < *P* < 0.975). The likelihood varied more erratically with changes in *a*, with two peaks and the second being within the 95% confidence interval of the maximum likelihood.

**Figure 3:**
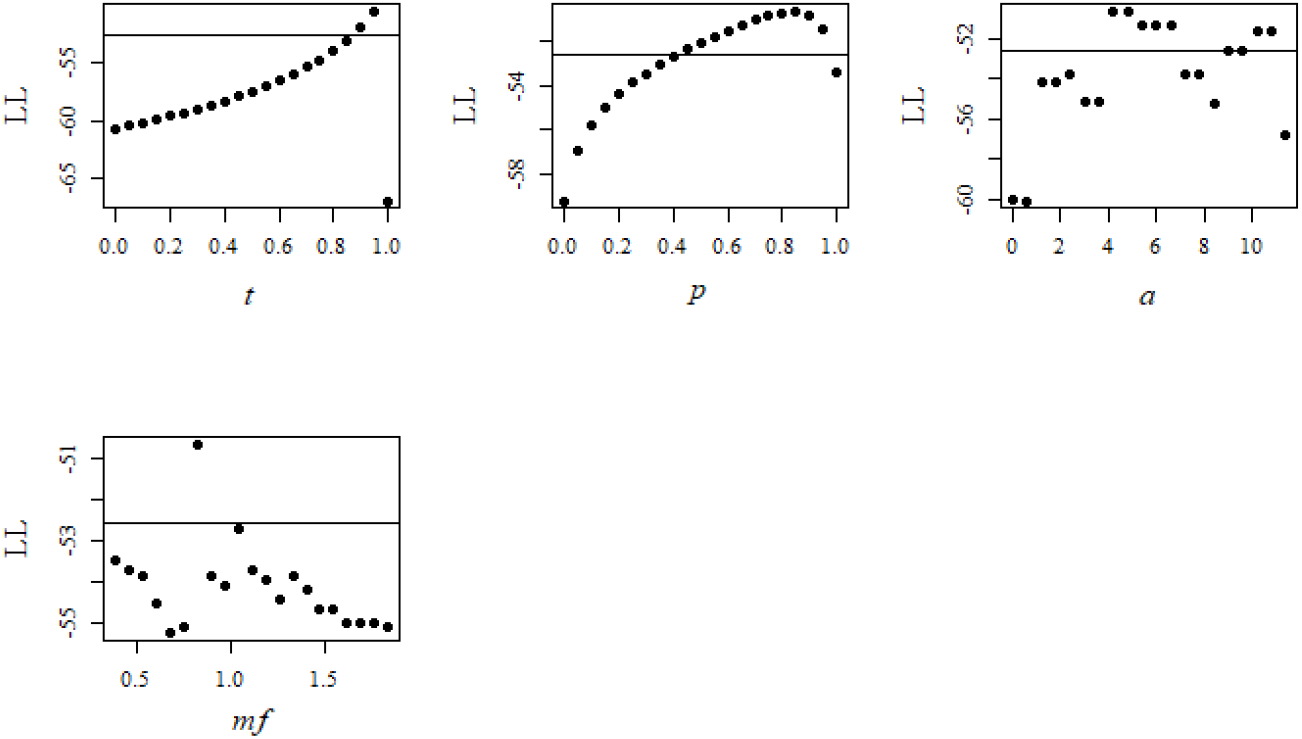
Variation in log-likelihood when *t, p, a* and *mf* are not fixed for model H15. The horizontal line is set at LL = -(negative log-likelihood + Chi-squared (*P* = 0.95, df =1) /2). This value of LL represents a significant difference in log-likelihood between the point under and over the line.

### Female attractiveness

The minimum adequate model (Table 3) showed that female attractiveness increased when female age, female pronotum width and male to female pronotum width ratio decreased. On the other hand, female attractiveness increased when male age increased. The decrease in female attractiveness in older females was stronger in larger females. The increase in female attractiveness when male age increased was weaker in larger males.

**Table 3:** Minimum adequate model (model *I*) predicting the number of attempts received by females

| Variables | Estimate | Standard error | *p*-value |
| --- | --- | --- | --- |
| Female age | - 0.4626 | 0.1612 | 0.0149 |
| Female pronotum width | - 0.9181 | 0.2961 | 0.0041 |
| Male age | 0.3508 | 0.0879 | 0.0019 |
| Male pronotum width | 1.0354 | 0.3487 | <0.0001 |
| Male to female pronotum width ratio | - 4.0798 | 1.3714 | 0.0029 |
| Interaction between female age and female pronotum width | 0.1379 | 0.0405 | 0.0006 |
| Interaction between male age and male pronotum width | - 0.0683 | 0.0277 | 0.0136 |

### Female receptivity

The minimum adequate model (Table 4) indicated that female receptivity increased when male age increased and when female pronotum width decreased. Female age, male pronotum width, the male to female pronotum width ratio, and the interactions between female age and female pronotum width and between male age and male pronotum width did not affect female receptivity.

**Table 4:** Minimum adequate model (model *J*) and non-significant terms predicting the number of matings by each female

| Variables | Estimate | Standard error | *p*-value |
| --- | --- | --- | --- |
| Female age | - 0.1135 | 0.0885 | 0.1999 |
| Female pronotum width | - 0.4548 | 0.2086 | 0.0292 |
| Male age | 0.1594 | 0.0560 | 0.0044 |
| Male pronotum width | - 0.1787 | 0.9225 | 0.8464 |
| Male to female pronotum width ratio | - 1.6523 | 1.1268 | 0.1425 |
| Interaction between female age and female pronotum width | 0.0728 | 0.11661 | 0.5319 |
| Interaction between male age and male pronotum width | - 0.0175 | 0.0994 | 0.860 |

## DISCUSSION

Similar to Caetano and Hajek (2017), we found that females are polyandrous with two thirds of the females that mated mating more than once. Like in other lek forming species, we also demonstrated a mating skew in males with only 28 of the 168 males mating (Widemo and Owens 1995). Interestingly, we found that only 19.7% of the contacts between a male and a female led to a mating attempt by the male. This suggests that males also exhibit choice and do not attempt to mate with all the females they encounter. Male mate choice has rarely been demonstrated in lekking insect species (Sæther et al. 2001; Werner and Lotem 2003; Shelly et al. 2012), but *S. noctilio* might be a new example. This should, however, be investigated and confirmed using bioassays where the same males are given access to multiple females at a time or in sequence.

The number of mating attempts made by males increase with male age (model I) or increases in males older than four days old (models G15 and G7). This increase in the number of attempts with male age could be explained by two phenomena that are not mutually exclusive. First, it is possible that males are sexually immature at emergence and need a few days to reach sexual maturity. This phenomenon has been observed in other hymenopteran species and is linked to delayed sperm maturation after emergence (Quimio and Walter 2000; Moors et al. 2009; Poidatz et al. 2018). Second, male *S. noctilio* might become more eager to mate as they age or might become less choosy with time (Bonduriansky 2001). This phenomenon has been observed in other insect species [e.g.

*Propylea dissecta* (Pervez et al. 2004), *Drosophila pseudoobscura* (Dhole and Pfennig 2014), *Drosophila melanogaster* (Dukas and Baxter 2014) *and Cimex lectularius* (Wang et al. 2016)]. To determine which of these two phenomena contributes to the effect of age on male motivation to mate in *S. noctilio*, sperm production and maturation in relation to age should be investigated.

Older males mated more (models H15, H14 and J). Similar to the effect of age on the number of attempts exhibited by males, the relationship between male age and mating success might be due to males needing to reach sexual maturity (Poidatz et al. 2018). Model J demonstrated that this increase in mating success in older males was not only linked to an increase in the number of attempts by males, but also to an increase in female receptivity as males aged. Females choosing to mate with older males is common among insects (Zuk 1987; Avent et al. 2008; Somashekar and Krishna 2011). It has been argued that male age gives females an estimate of male fitness because, by surviving longer, older males demonstrate genetic superiority (Manning 1985; Brooks and Kemp 2001). This hypothesis is particularly relevant considering that males emerge a few days before females. While they are waiting for females to emerge, males are exposed to predators and fluctuating environmental conditions that can significantly reduce their lifespan and prevent them from reproducing.

While *H15* indicated that mating success increased in males older than four days old, the distribution of the likelihood of *a* within model H14 was bimodal, with a peak around four days and a peak around 10 days. This suggested that the increase in mating success is equally likely in males older than four days old and in males older than ten days old. The fact that these two models did not reach a consensus could indicate that time is only one of the factors influencing sexual maturity in males. For example, insect development and maturation are very sensitive to temperature (Rockstein and Miquel 1973). While we tried to keep temperature constant where the wasps were stored, it is possible that individuals being born and aging on slightly warmer days could have aged faster and reached sexual maturity sooner (Jaycox 1961; Akman Gündüz and Gülel 2002). Both environmental and individual variation could lead to differences in the time to sexual maturity in males. As a result, although we observe a strong effect of age on mating success, it is impossible to give an exact estimate of the age at which *S. noctilio* males exhibit the highest mating success.

Female attractiveness decreased as females aged and this effect was stronger in larger females (model I). This observation can be explained by two non-mutually exclusive phenomena. First, males might prefer mating with younger females, like in *Ephestia kuehniella* (Lepidoptera: Pyralidae) (Xu and Wang 2009) or *Empis borealis* (Diptera: Empididae) (Svensson et al. 1989). Second, males might not recognize older females as potential mates. Böröczky et al. (2009) demonstrated the presence of a contact pheromone on the cuticle of the females in *S. noctilio* that triggers mating attempts from males. Perhaps older females have reduced titers of cuticular hydrocarbons and the contact pheromone. Indeed, in the Hymenoptera, females can produce male progeny without being fertilized. After reaching a certain age, whether they are mated or not, *S. noctilio* females switch their focus from mate location to host location for egg laying (Morgan and Stewart 1966). This succession of behaviours is similar to other hymenopteran species in which mates and hosts are separated in the landscape (Guertin et al. 1996). The mechanisms that trigger a switch from a mate-searching to a host-searching phase in unmated hymenopteran females are still unknown, but it is likely that this switch would be characterized by a change in behaviour and physiology. In *S. noctilio*, those physiological changes might include qualitative and/or quantitative changes in the cuticular hydrocarbon profile of females. The relationship between contact pheromone production and female age should be investigated as it could greatly influence the success of laboratory rearing programs.

It is interesting to note that female age did not influence female receptivity. Because *S. noctilio* is haplodiploid, unmated females can still produce sons. As a result, the pressure to mate might not be as strong as in diploid species. Furthermore, if female age did not influence female receptivity but influenced female attractiveness, it would suggest that males can recognize receptive females. Indeed, female receptivity was determined as the proportion of attempts from males accepted by females. This means that female receptivity could not be evaluated in non-attractive females. The hypothesis that males can detect receptive females agrees with the hypothesis that female behaviour and physiology are synchronized and change from a mate-locating to a host-locating phase (Morgan and Stewart 1966).

Larger males attempted to mate more than smaller males (model I). Larger males are likely to carry more sperm that smaller males (Schlüns et al. 2003; Pech-May et al. 2012). It is possible that those males are less picky and would try to mate with females that smaller males would consider unsuitable (Byrne and Rice 2006).

It is interesting to note that models G15 and G7 did not find any effect of male size on mating attempts, except when male size was compared to female size. Models G15 and G7 suggested that males smaller than 103% the size of the females attempted to mate more than larger males. Similarly, model I suggested that female attractiveness increased when the male to female size ratio decreased. This indicates that regardless of their size, males attempt to mate with females that are larger than them or similar in size. During the bioassays, we observed that large males that tried to mate with small females had trouble bending their abdomen far enough, preventing them from fertilizing these females. This phenomenon was also observed by Caetano and Hajek (2017).

The observation that males prefer mating with females larger than them is also reflected in the effect of the size ratio on mating success. Model H15 indicated that males that were smaller than the female (i.e. 82% of the size of the female) had the highest mating success.

Caetano and Hajek (2017) also tested the effect of male to female size ratio on the mating success of *S. noctilio*. Contrary to our results, Caetano and Hajek (2017) did not find any effect of size on mating success. However, their study did not include age as an explanatory variable. In our study, male and female age seemed to be important factors explaining mating success, and not including these variables as predicting factors could have concealed the effect of size.

It is important to note that the effects of male size and size ratio on mating success observed in our bioassays, might be absent in the field. While larger males might struggle to fertilize small females, it is possible that larger males benefit from their size in different ways. During the bioassays, we often observed aggressive interactions between male *S. noctilio*. In *Polistes dominulus* (Hymenoptera: Vespidae) males also exhibit strong aggression and compete for territories within the lek (Izzo and Tibbetts 2012). After a fight, the winning male usually has a higher mating success. Male size could be a determining factor in the outcome of male-male contests and play an important role in the establishment of a hierarchy within the lek. It is possible that bioassays in captivity do not include some size-dependent dynamics that are present in the field. In the future, the timeline of lek formation and hierarchy establishment within leks of *S. noctilio* should be investigated. Large body size could also be associated with larger amounts of sperm (Schlüns et al. 2003; Pech-May et al. 2012). Larger males might be able to fertilize a greater number of females or to offer more sperm during fertilization.

Female receptivity increased as female size decreased (model J). This result seems counter intuitive because smaller females produce fewer eggs (Morgan and Stewart 1966). Small females might be unable to reject males and accept mating to avoid harassment and injuries. Convenience polyandry (Thornhill and Alcock 1983), or females accepting mating because they cannot reject males, is frequently observed in insects, including the Hymenoptera (Fernández-Escudero et al. 2002; Trontti et al. 2007). Model I also suggested that small females are more attractive to males. This echoes the convenience polyandry hypothesis because it might be easier for males to force small females into mating.

## Conclusions

The results of this study suggested that over their lifetime, the mating success of male and female *S. noctilio* is dictated by the motivation of each sex to mate. Males might need to reach sexual maturity to mate while older females probably switch their behaviour from mate to host location. When both sexes are willing to mate, females prefer older males. Small males seem to also be at an advantage for fertilization because they can fertilize small females. However, it is still unclear if females use other male traits to select males in natural conditions. These traits could be related to genetic compatibility (Thiel et al. 2013), male motor performance (Byers et al. 2010) or the male hierarchy within the leks (Izzo and Tibbetts 2012). *Sirex noctilio* exhibit sexual dimorphism with males having an orange abdomen while the females are fully black. The fact that the presence of dead male *S. noctilio* increases the capture of females by unbaited traps (Allison et al. 2019) suggests that the orange colouration of the males also plays a role in mate choice. Each of these hypotheses deserve investigation in laboratory and field conditions. Finally, our results suggest the presence of mate choice by male *S. noctilio*. This type of behaviour is rarely investigated in lekking species (Sæther et al. 2001; Werner and Lotem 2003; Shelly et al. 2012). For this reason, mate choice by males should receive more attention in *S. noctilio*. Our results suggest that males do not select females based on size. Perhaps other cues or signals such as genetic compatibility, or sex pheromones are used by males to evaluate females.

The future development of new management strategies might benefit from our findings. For example, gene drive technologies and sterile insect techniques rely heavily on the capacity of the lab-reared males to be as attractive to females as wild males. Our results suggest that males younger than 4 days old are not sexually mature. Therefore, releasing immature males could compromise the success of a mass release. Similarly, ignoring the presence of age-dependent behaviours and physiological states might have compromised the discovery and study of potential sexual pheromones produced by *S. noctilio*.

## Acknoledgements

We thank members of the Tree Protection Cooperative Programme (TPCP), the Department of Agriculture, Forestry and Fisheries (DAFF), the National Research Foundation (NRF) of South Africa, Natural Resources Canada and the USDA-FS FHP for funding. We also thank members of the TPCP and the South African Sirex Control Programme for assistance with field work and sample collection.

